# A CREBZF/Zhangfei isoform activates CHOP during prolonged cellular stress

**DOI:** 10.1101/2022.10.10.511668

**Authors:** Yu Li, Wan Kong Yip, Jenna Penney, Tiegh Taylor, Yani Zhang, Minghua Zeng, Timothy E. Audas, Ray Lu

## Abstract

The basic leucine zipper transcription factor CREBZF (Zhangfei or ZF) was identified through its interaction with Herpes Simplex Virus-1 related cellular protein HCF-1. CREBZF has been implicated in cellular stress responses through its interaction with other proteins, such as CREB3/Luman and ATF4. Here we investigated the production of four CREBZF isoforms, which arise from translational initiation of a downstream AUG at codon 83, and mRNA alternative splicing that adds an IFFFR pentapeptidyl tail to the C-terminus. We found that in addition to transcriptional activation, the short-tailed CREBZF (stZF) isoform was specifically induced by prolonged ER stress treatment. This stZF isoform is a potent transcriptional activator of the pro-apoptotic protein CHOP. Overexpression of stZF activates transcription of CHOP through a CCAAT enhancer binding protein (C/EBP)-ATF site, and promotes apoptosis. We propose that 1) CREBZF is a key component of the Integrated Stress Response (ISR); 2) stZF is essential for the role of CREBZF in inducing CHOP and promoting cell death upon prolonged cellular stress.

## INTRODUCTION

Perturbing homeostasis of the endoplasmic reticulum (ER) triggers the unfolded protein response (UPR). The UPR is largely an adaptive response that includes attenuating protein synthesis, enhancing protein folding capacity, and increasing protein degradation (Bailey and O’Hare, 2007; Rutkowski and Kaufman, 2004; Schroder and Kaufman, 2005; Yoshida, 2007). There are three ER-resident transmembrane proteins identified as major proximal sensors in mammalian cells that respond to ER stress: protein kinase-like endoplasmic reticulum kinase (PERK), activating transcription factor 6 (ATF6), and endoribonuclease inositol-requiring enzyme-1 (IRE1) (Haze et al., 1999; Liu et al., 2000; Rutkowski and Kaufman, 2004; Schroder and Kaufman, 2005). When all the measures to alleviate ER stress fail, the cell switches to a pro-apoptotic response and leads to cell death.

An important ER stress-associated apoptosis signaling pathways is mediated by C/EBP homologous protein (CHOP), also known as growth arrest- and DNA damage-inducible gene 153 (GADD153) (Ma et al., 2002; Rutkowski and Kaufman, 2004; Zinszner et al., 1998). CHOP represses transcription of the anti-apoptotic gene Bcl-2, thereby shifting the balance towards the pro-apoptotic family members (Bad, Bak, and Bax) (McCullough et al., 2001; Schroder and Kaufman, 2005). CHOP is known to be transcriptionally regulated by ATF4 and ATF6, two key sensors of the UPR. ATF4 can activate CHOP via a C/EBP-ATF composite element in the CHOP promoter which responds to ER stress as well as amino acid deprivation (Bruhat et al., 2000), while ATF6 stimulates CHOP transcription through an ER-stress response element (ERSE) (Oyadomari and Mori, 2004; Yoshida et al., 2000).

CREBZF was identified as Zhangfei (ZF), a basic region leucine zipper (bZIP) transcription factor that interacts with the Herpes Simplex Virus-1 (HSV-1) related host cellular protein HCF-1(Kristie et al., 1995; LaBoissiere and O’Hare, 2000; Wilson et al., 1993), and inhibits HSV-1 replication when overexpressed in cell cultures (Akhova et al., 2005; Lu and Misra, 2000b). Like pro-apoptotic CHOP, CREBZF is unable to bind DNA as a homodimer (Hogan et al., 2006; Lu and Misra, 2000b). Compared to the DNA-binding basic regions of other bZIP proteins, CREBZF has an unusual amino acid sequence and a different predicted protein structure (Cockram et al., 2006). Besides its potential role in HSV-1 replication (Lu and Misra, 2000b), CREBZF has been shown to mediate medulloblastoma cell differentiation and apoptosis by inducing expression of the nerve growth factor receptor trkA (Valderrama et al., 2009), and cell survival and growth via p53 (Bodnarchuk et al., 2012; Zhang and Misra, 2014). CREBZF can also act as a co-regulator of nuclear receptors in cell type-specific gene expression (Xie et al., 2008; Xie et al., 2009a; Xie et al., 2009b). CREBZF associates with SHP (small heterodimer partner) and inhibits gene expression mediated by nuclear receptors such as estrogen and glucocorticoid receptors (Xie et al., 2008; Xie et al., 2009b). In regards to the UPR, we have previously found that CREBZF can function as a coactivator of ATF4 (Hogan et al., 2006), although it represses the transactivation activity of Luman/CREB3 (Misra et al., 2005).

The original 272-amino acid Zhangfei/ZF was considered the predominant form of CREBZF protein (Akhova et al., 2005; Cockram et al., 2006; Hogan et al., 2006; Lu and Misra, 2000b; Misra et al., 2005). With revisions of the human genome sequence, it was recently reported that a longer isoform is the most common form detected in cell lines and mouse tissues, which starts from an upstream in-frame AUG and adds an extra 82-amino acids to the N-terminus (Xie et al., 2008).

Here we report that, in addition to the long and the short isoforms (lZF and sZF), CREBZF can also produce two other isoforms by alternative splicing of a conserved 213-base region of the mRNA, resulting in the removal of only the original stop codon and the addition of a pentapeptide tail, IFFFR, to the C terminus. We found that only the short-tailed CREBZF isoform (stZF) could potently activate CHOP gene expression. We mapped the stZF response element to the C/EBP-ATF composite site in the CHOP promoter (Bruhat et al., 2002; Bruhat et al., 2000). Furthermore, prolonged treatment with ER stressor specifically induced stZF, and overexpression of the tailed isoforms promoted cell death. Since mRNAs of the spliced CREBZF isoforms are thought to be targets of nonsense-mediated mRNA decay (NMD) (reviews Matsuda et al., 2008; Stalder and Muhlemann, 2008), we postulate that during prolonged UPR, the cell may stabilize the spliced mRNA variants to shift the survival/death balance towards apoptosis.

## RESULTS

### CREBZF can produce four isoforms through alternative splicing and differential translational initiation

The GenBank entry of CREBZF (GeneID: 58487) (at the time of this manuscript submission) shows that the transcript is an unspliced mRNA (Fig. 1A, variant 1), which encodes a polypeptide of 354 amino acids. In addition, there are four other transcript variants listed as non-coding because they are substrates of nonsense-mediated mRNA decay (NMD). NMD is an mRNA surveillance system that is conserved in all eukaryotes to detect and degrade mRNAs containing premature translation termination codons (Matsuda et al., 2008; Stalder and Muhlemann, 2008). Mammalian cells are estimated to use NMD to downregulate approximately 1/3 of alternatively spliced mRNAs, most of which are believed to be erroneous mRNA splicing products (Lareau et al., 2004; Lewis et al., 2003). Of the 4 transcripts of NMD substrate, two are shown in Fig. 1A as variants ***b*** and ***c***; the remaining two variants (not shown) lack one or two more internal segments in nonprotein coding exon 3 of variant ***c***. All these splicing variants are NMD substrates because they meet the “50- to 55-nucleotide rule”, which stipulates that translation termination at a nonsense codon located > ~50 to 55-nucleotides upstream of an exon-exon junction and triggers NMD (Matsuda et al., 2008; Stalder and Muhlemann, 2008). To date, only a few genes have escaped from this rule (Buhler et al., 2004; Carter et al., 1996; Inacio et al., 2004; Kim et al., 1998; Zhang and Maquat, 1997). CREBZF mRNA splicing variants also have an unusually long 3’-untranslated region (UTR) that is known to promote NMD (Matsuda et al., 2008; Stalder and Muhlemann, 2008).

**Figure 1.**
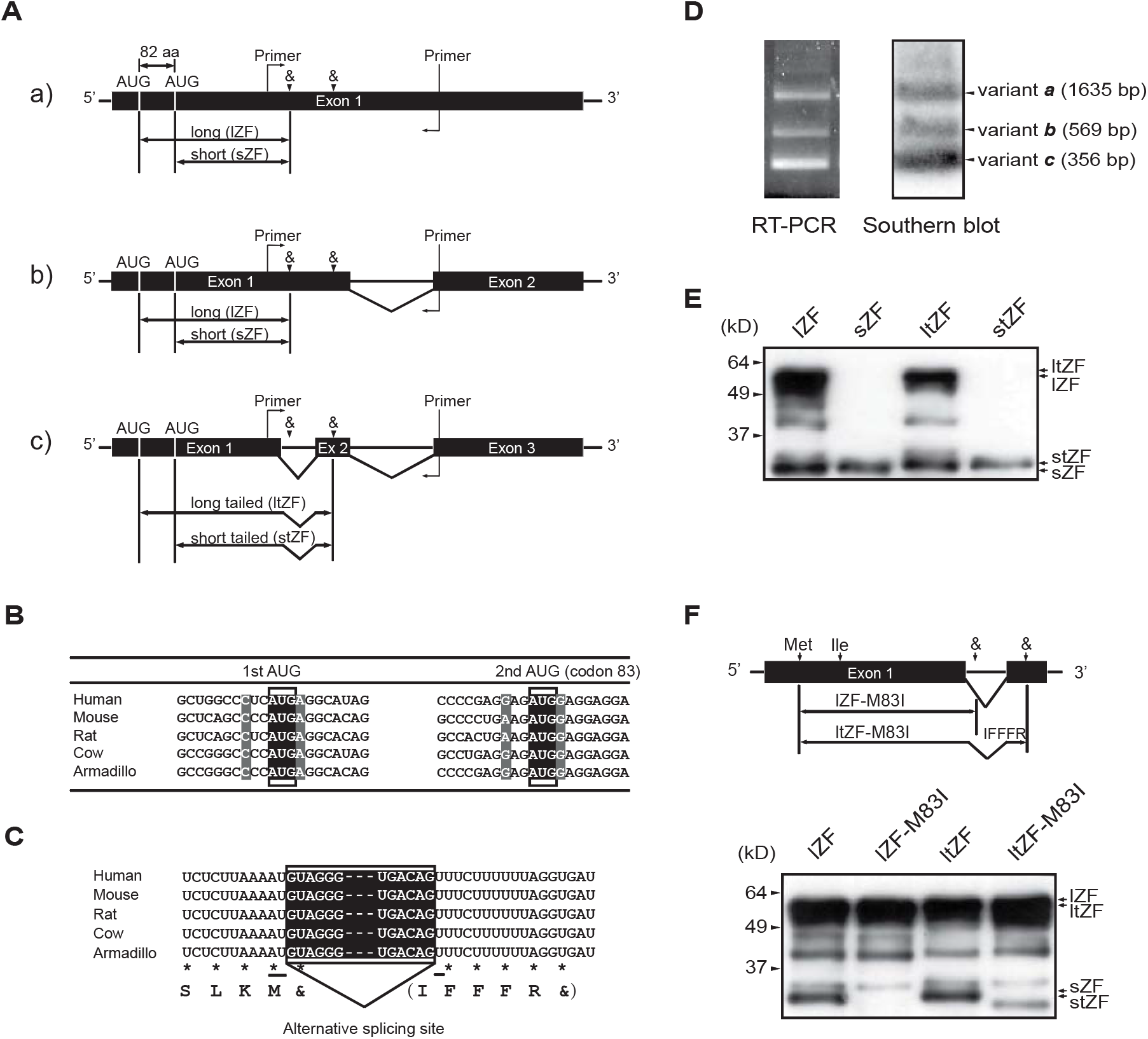
The human CREBZF gene can produce four isoforms. (A) Schematic representation of human CREBZF mRNA splicing variants. Alternative use of the two AUGs as translational initiation sites produces the long (lZF) and the short CREBZF protein (sZF); with an additional mRNA splicing event found in variant ***c***, two more isoforms can also be produced: the long CREBZF with a C-terminal tail (ltZF) and the short polypeptide with the tail (stZF). “&” denotes a stop codon. (B) Alignment of the mRNA sequences of CREBZF homologs adjacent to the two translational initiation sites. The bases at positions −3 and +4 relative to the initiation codon are shaded grey which are important for the Kozak sequence. (C) Alignment of CREBZF homolog mRNA sequences at the splicing junction of the first intron in Splicing variant ***c***. The bases in the shaded box are the 213-nucleotide intronic sequence. This alternative splicing event removes the stop codon of the original transcript and leads to the addition of a pentapeptide, IFFFR, tail to the C-terminus of the protein. (D) Detection of the splicing variants. Total RNA was prepared from HeLa cells, and reverse transcriptase PCR (RT-PCR) was carried out using primers flanking the alternatively spliced region (left panel). The RT-PCR fragments were verified by Southern blotting using a CREBZF cDNA probe (right panel). (E) Production of the four CREBZF protein isoforms. HeLa cells were transfected with cDNAs encoding lZF, sZF, ltZF or stZF. Western blotting was performed using anti-CREBZF (Rb6) antibody. (F) Mutation of the AUG at codon 83 to isoleucine eliminates the production of short CREBZF isoforms by lZF and ltZF cDNA constructs. HeLa cells were transfected with cDNAs encoding lZF, ltZF or mutants lZF-M83I and ltZF-M83I. Western blotting was carried out using the anti-CREBZF (Rb6) antibody.

Using the online transcript mapping tool, Aceview (http://www.ncbi.nlm.nih.gov/IEB/Research/Acembly/) (Thierry-Mieg and Thierry-Mieg, 2006), we found that, although the unspliced variant ***a*** had the most (a total of 341) supporting cDNA clones, variant ***c*** had 177 supporting cDNA clones, which was significantly more than variant ***b*** (49 clones) or the combined total of the remaining variants (16 clones). We believe that this high prevalence implies that the cell may specifically stabilize variant ***c*** under certain conditions. Although not coding for proteins, the nucleotide sequence in the junction region between exon 1 and exon 2 was found to be highly conserved, implicating an important function of these sequences (Fig. 1C). Remarkably, this alternative splicing precisely skips the original stop codon, but only adds a pentapeptide tail, IFFFR, to its C-terminus (Fig. 1C).

We had believed that translation of CREBZF mRNA starts preferentially from the second AUG, because of the existence of a consensus Kozak sequence (a purine at −3 and a G at +4 position) (Kozak, 1997, 2005) around the second AUG and a would-be “classic” N-terminal activation domain following codon 83 (Cockram et al., 2006; Hogan et al., 2006; Lu and Misra, 2000b) (Fig. 1B). More recent data seem to suggest that the predominant or default translational start is from the first AUG (Xie et al., 2008).

Current data suggest that CREBZF is a protein of multiple functions. To investigate the potential association between its different functions with specific isoforms, we sought to identify the isoforms that CREBZF might produce. Reverse transcriptase PCR (RT-PCR) of the total RNA extracted from HeLa cells was carried out by using PCR primers flanking the intron and alternative splicing site (Fig. 1A). This resulted in the amplification of three distinct DNA fragments, which agreed perfectly with the predicted sizes (1635, 569 and 356 bp) of the three transcript variants (Fig. 1D). Southern blotting and DNA sequencing were also used to confirm the identities of these bands.

To examine whether the second AUG at codon 83 can indeed be used for translational initiation, we transfected HeLa cells with plasmids containing cDNAs corresponding to lZF, sZF, ltZF and stZF, and checked their protein products by Western blotting. The cDNAs of sZF and stZF produced proteins with apparent molecular mass of ~29 kDa; in addition to these bands, their long CREBZF counterparts (lZF and ltZF) produced bands of ~52 kDa (Fig. 1E). These results suggest that the AUG at codon 83 can also be used as translational initiation.

To rule out the possibility that the presumed sZF and stZF bands (Fig. 1E, lane 1 and 3) were degradation products of the long CREBZF proteins, we mutated the second ATG to ATC (an isoleucine codon) in the cDNA and generated lZF-M83I and ltZF-M83I mutants (Fig. 1F). Transfection of these mutant cDNAs into HeLa cells resulted in the elimination of the 29-kDa polypeptides, leaving only the 52 kDa bands. These results indicated that the AUG at codon 83 can be an alternative translation initiation site, probably through a leaky scanning mechanism of the ribosome.

We noticed the actual molecular mass (~52 kDa) of lZF and ltZF were much greater than the calculated 37 and 38 kDa. The significantly larger actual molecular mass of long CREBZF isoforms may be due to post-translational modifications on the extra 82 amino acids at the N-terminus.

### ER stress induces short CREBZF with the pentapeptide tail (stZF) and overexpression of stZF increases protein level of CHOP

We have recently discovered that the transcription of CREBZF is induced by amino acid deprivation (Zhang et al., 2010). Since previously we were unable to detect changes of CREBZF expression in cells under common ER stress treatments, we thus asked whether prolonged ER stress would induce certain CREBZF isoforms. We treated RAW264.7 (mouse macrophage) cells, which has the highest endogenous CREBZF among the cell lines that we studied, with tunicamycin, an ER stressor that blocks N-linked glycosylation, and analyzed CREBZF expression by Western blotting (Fig. 2A). Consistent with this notion, endogenous stZF was induced RAW264.7 after 24 and 48 hours of stress treatment. We further explored whether overexpression of stZF affects expression of proteins associated with the UPR. RAW264.7 cells were transfected with pcDNA3.1 or stZF, followed by stress treatment with 5 μg/ml tunicamycin. Total cell lysates were analyzed by Western blotting (Fig. 2B). The levels of chaperone BiP/GRP78 (Kohno et al., 1993) were used as a positive indicator of ER stress in the cells. Protein level of CHOP was increased in cells overexpressing stZF at 17- and 24-hr time points.

**Figure 2.**
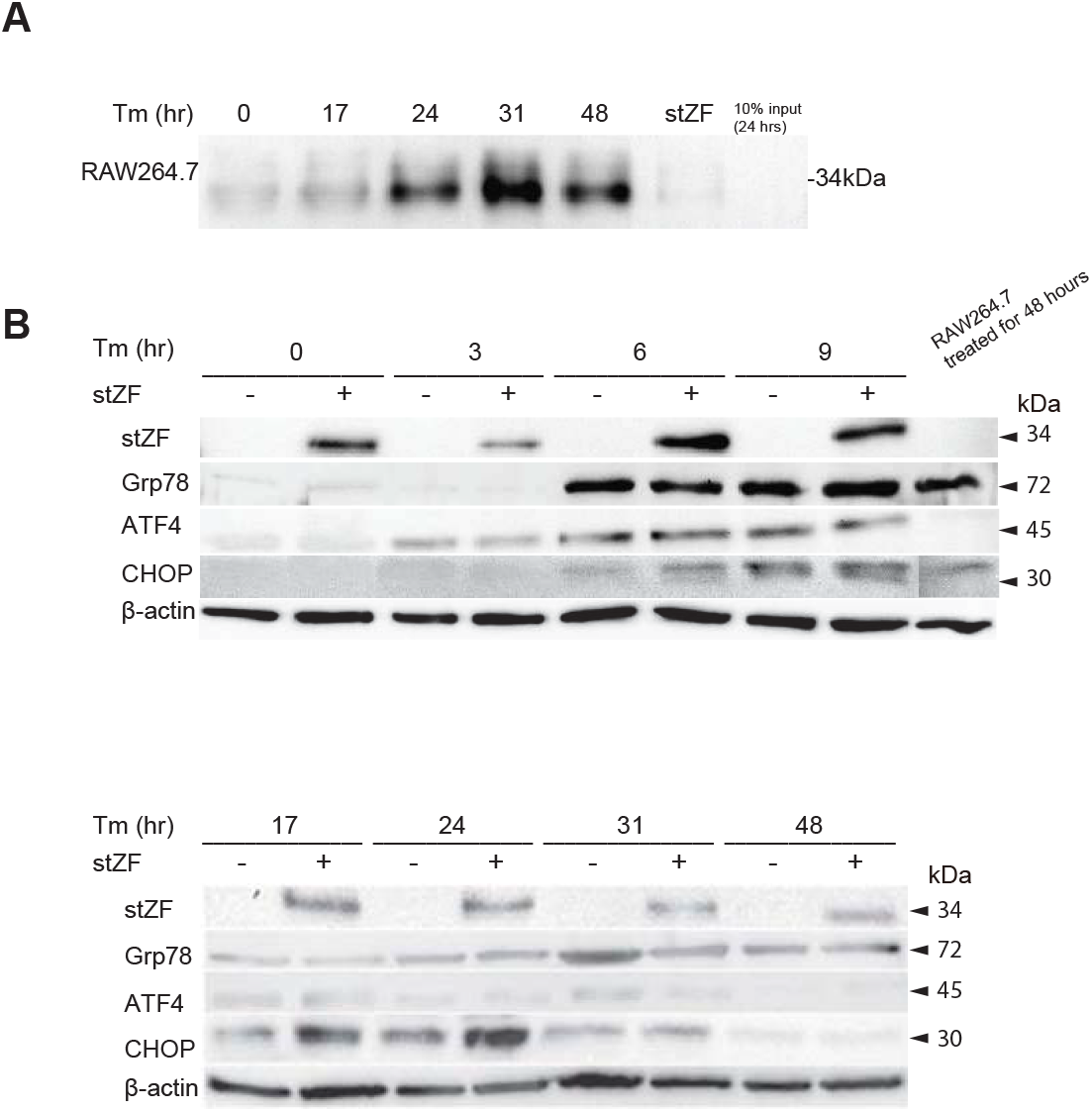
stZF is induced by prolonged ER stress and overexpression of stZF increases protein level of CHOP. (A) RAW264.7 cells with 5 μg/ml tunicamycin for indicated length of time. Cell lysates were immunoprecipated prior to western blotting. Cells transfected with stZF were used as a protein size marker. (B) RAW264.7 cells were transfected with pcDNA3.1 or stZF and treated with 5 μg/ml tunicamycin. Overexpression of stZF increased protein level of CHOP after 17 and 24 hours of exposure to the ER stressor. Expressions of Grp78 and ATF4 were analyzed, but their protein levels were not affected by stZF overexpression.

### CHOP is a downstream target of stZF and C/EBP-ATF is the response element of stZF in the CHOP promoter

Since sZF is a transcription factor with a potent N-terminal activation domain (Cockram et al., 2006; Hogan et al., 2006; Lu and Misra, 2000b) and overexpression of stZF increased protein level of CHOP (Fig. 2B), we sought to investigate whether stZF regulates CHOP at the transcriptional level. Dual luciferase assays were conducted using a firefly reporter plasmid containing a CHOP promoter fragment (−954 to +1). In agreement with the Western blot results, while lZF and sZF had no effect on the transcriptional activation of the CHOP promoter, CREBZF isoforms containing the pentapeptide IFFFR tail stimulated transcription of the CHOP promoter, with stZF sample having the highest level of activation (2.8 fold) (Fig. 3A).

**Figure 3.**
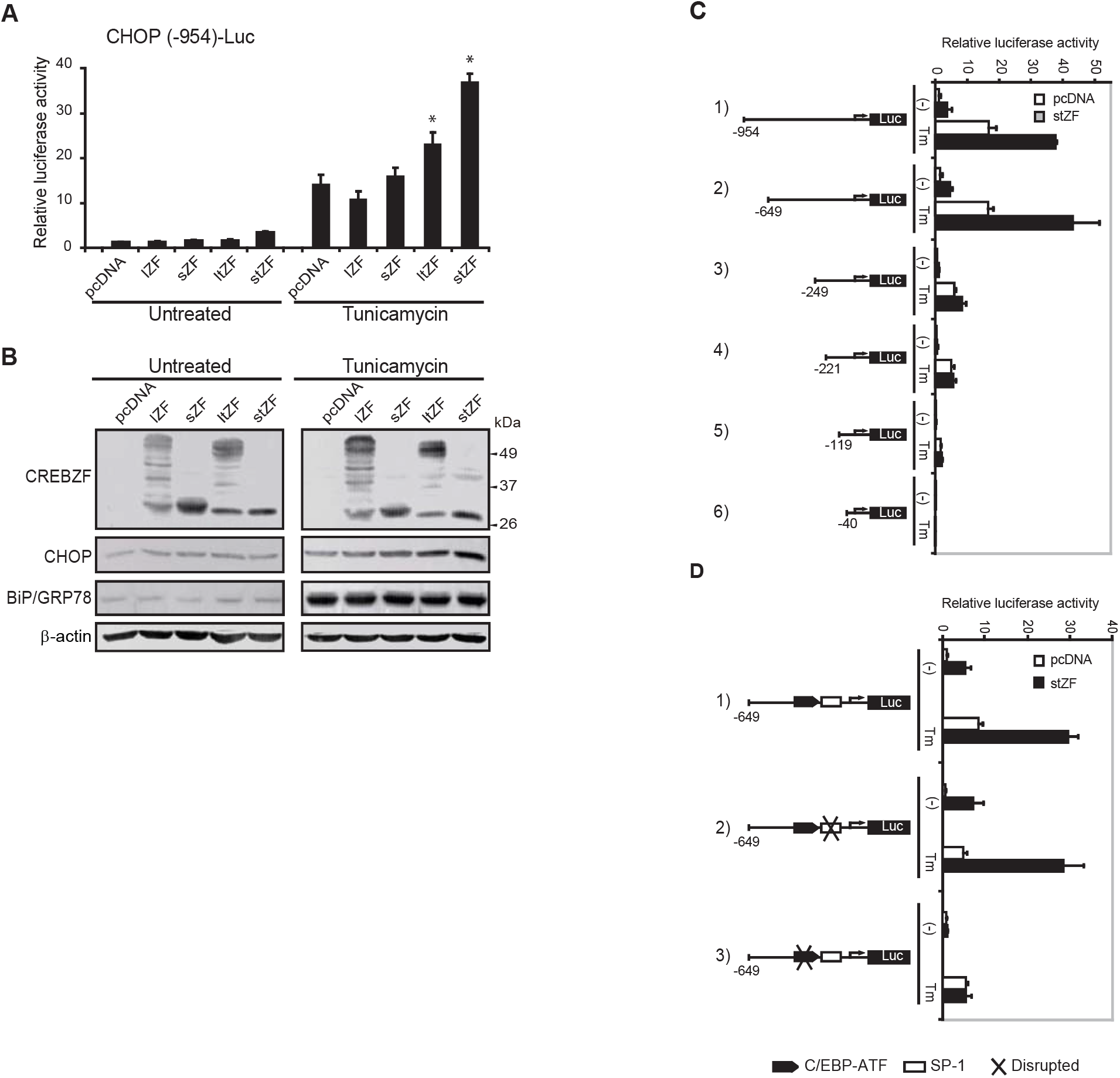
stZF induces expression of the pro-apoptotic gene CHOP and mapping of a stZF response element in the CHOP promoter. (A) CHOP promoter reporter assay. HeLa cells were transfected with a luciferase reporter containing the CHOP promoter between nucleotide position −954 and +91 together with the reference plasmid pRL-SV40 and cDNAs encoding lZF, sZF, ltZF or stZF. Cells were treated with tunicamycin (black bars) as described above and dual luciferase activities were measured. (B) Western blot analysis of HeLa cells transfected with pcDNA3.1 or cDNAs encoding lZF, sZF, ltZF or stZF. Twenty four hours post-transfection, cells were treated with 2 μg/ml tunicamycin for 24 hours. (C) Deletion mapping of the CHOP promoter. HeLa cells were transfected with luciferase reporters containing progressive 5’ deletions of the CHOP promoter together with the reference plasmid pRL-SV40 and cDNA encoding stZF. (D) The C/EBP-ATF element is essential for CHOP promoter activation by stZF. HeLa cells were co-transfected with luciferase reporters containing the CHOP promoter DNA fragment (−649/+91) or the mutant containing a disrupted SP-1 or C/EBP-ATF element together with the empty plasmid pcDNA3.1 or cDNA encoding stZF. The position of the SP-1 and the C/EBP-ATF composite site are as indicated. Twenty four hours post-transfection, cells were treated with 2 μg/ml tunicamycin for 24 hours prior to determination of luciferase activities.

We transfected HeLa cells with the four CREBZF isoforms and analyzed the level of CHOP protein by Western blotting (Fig. 3B). We found that the CHOP protein was induced by overexpression of ltZF and stZF, especially in response to 2 μg/ml tunicamycin treatment for 24 hrs (Fig. 3B, the last two lanes in the right panel), with stZF being the strongest inducer. Next, we decided to map the response element of stZF in the CHOP promoter by deletion analysis. We used progressive 5’ deletions of the CHOP promoter and performed reporter assays (Fig. 3C) in stZF-transfected cells with or without tunicamycin (Tm) treatment. We found that deletion of the region −954 to −649 had no effect on transcriptional activation in both untreated and tunicamycin treated cells (Fig. 3C, row 2). However, deletion of the region between −649 and −249 reduced transcriptional activation most significantly (by 70%), while subsequent deletions had no obvious effect. These results suggest that the region between −649 and −249 contains a *cis*-element(s) that mediate transcriptional activation by stZF.

It has been reported that the region between −649 and −249 contains two response elements: a C/EBP-ATF composite site (ATTGCATCAT) and a SP-1 binding site (CGCCCC) (Bruhat et al., 2000; Wolfgang et al., 1997). To further map the stZF response element within the CHOP promoter, we mutated both of these sites on the CHOP promoter and performed luciferase reporter assays. We found that mutation of the C/EBP-ATF element abolished transcriptional activation of the CHOP promoter by stZF (Fig. 3D), while mutation of the SP-1 element had minimal effect (Fig. 3D). These results indicate that the C/EBP-ATF element is required for the CHOP promoter activation by stZF.

To ascertain whether the C/EBP-ATF element is sufficient for stZF-mediated transcriptional activation and also to compare the transactivation potential on this element by other CREBZF isoforms, luciferase assays using a 2×C/EBP-ATF-containing reporter were conducted. We found that transient transfection of stZF produced the most (~5-fold) activation of transcription compared to that of pcDNA3.1, while lZF and sZF had no effect (Fig. 4A). Western blot analysis of the transfected cell lysates indicated that the differences of transactivation potential were not due to the levels of the proteins.

**Figure 4.**
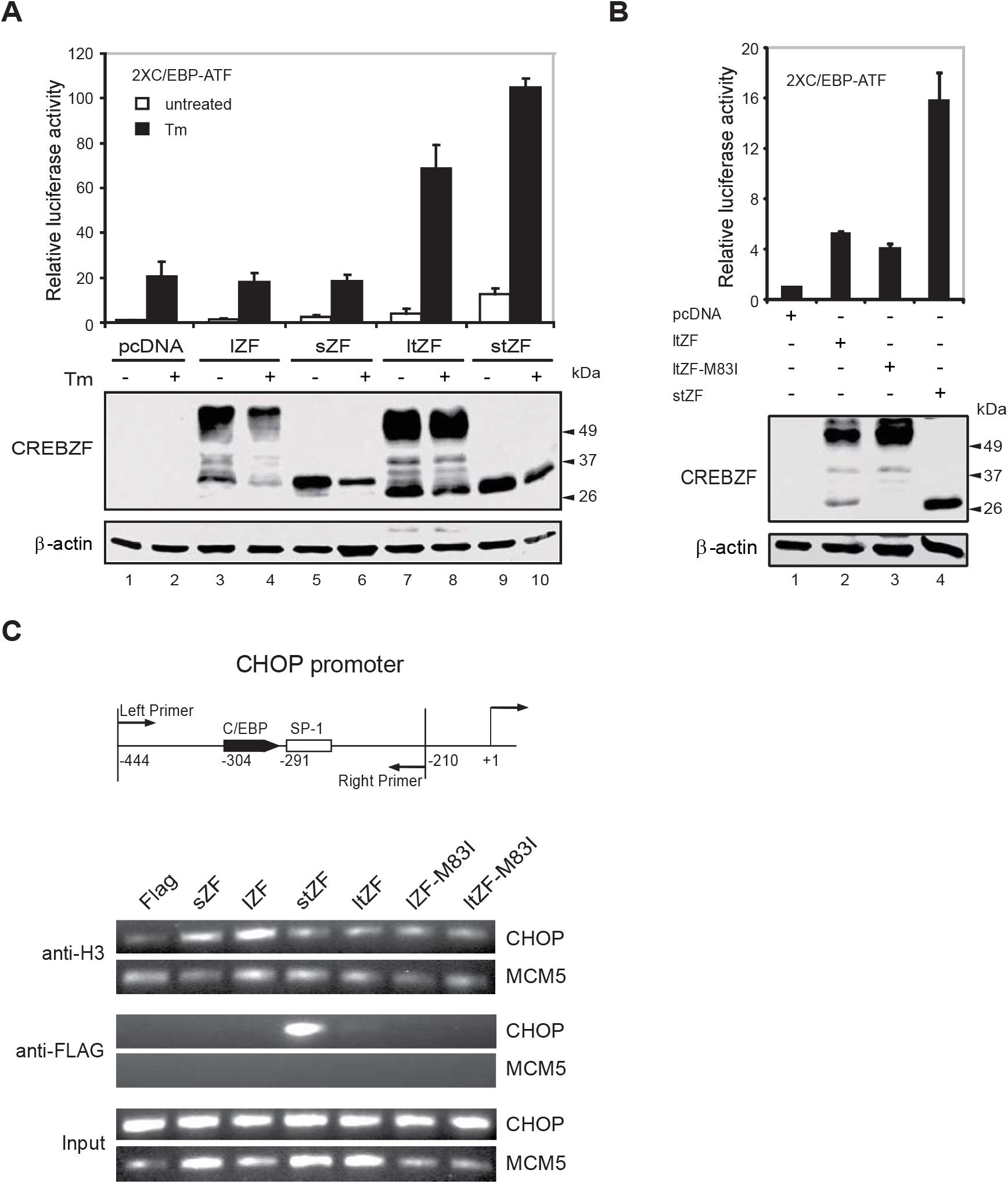
stZF binds and activates transcription from the C/EBP-ATF element in the CHOP promoter. (A) stZF activates transcription from a C/EBP-ATF reporter plasmid. HeLa cells were co-transfected with a luciferase reporter containing two copies of the C/EBP-ATF element together with the vector plasmid pcDNA3.1, or cDNAs encoding the four CREBZF isoforms, under untreated or tunicamycin-treated conditions. Samples were analyzed by dual luciferase assays and Western blotting as described above. (B) The stZF isoform is a potent transcriptional activator on the C/EBP-ATF containing promoter. HeLa cells were cotransfected with the 2×C/EBP-ATF reporter plasmid, as well as cDNAs encoding pcDNA3.1, ltZF, ltZF-M83I, and stZF. Forty eight hours after transfection, luciferase activities were measured, and the cell lysates were analyzed by Western blotting. (C) Direct binding of stZF to the CHOP promoter as measured by the ChIP assay. Top panel is a schematic diagram of the human CHOP promoter, with positions of the primer pair used in the ChIP assay indicated. HeLa cells were cotransfected with the pcDNA 3.1-based pcFLAG empty vector, or with cDNAs encoding the four isoforms, mutant lZF(M83I) or ltZF(M83I) along with the CHOP promoter, and then cross-linked with formaldehyde. Chromatin was immunoprecipitated with the indicated antibodies. Purified precipitates or input DNA was analyzed by PCR using primers specific for CHOP (−444/−210) or the control MCM5 promoter. PCR products was subjected to gel electrophoresis and visualized by ethidium bromide staining.

We noticed that ltZF also produced a 3.5-fold activation of the 2×C/EBP-ATF reporter (Fig. 4A). Since we knew that ltZF could produce short isoform stZF (Fig. 1F), it is possible that this observed activation was due to the stZF polypeptide produced by alternative translational initiation. To test this hypothesis, we used the mutant construct ltZF-M83I with the second ATG deleted, and performed the luciferase assays. We found that the activation potential of the ltZF-M83I mutant was indeed lower than that of ltZF, especially when the reporter activities are normalized against the protein levels (Fig. 4B).

We also performed chromatin immunoprecipitation assays to assess whether stZF directly binds to the CHOP promoter *in vivo* (Fig. 4C). HeLa cells were transiently transfected with plasmids expressing CREBZF isoforms fused with a FLAG epitope along with the CHOP promoter. After cross-linking and immunoprecipitation using specific antibodies as indicated, PCR was performed to detect the presence of the CHOP promoter region by using primers flanking the C/EBP-ATF element site (Fig. 4C, top panel). MCM5 was arbitrarily chosen as a negative control. We found that FLAG antibody readily precipitated chromatin containing the CHOP promoter specifically in the FLAG-stZF-transfected cells and weakly in the FLAG-ltZF cells, but not in the lZF or sZF isoforms (Fig. 4C). Taken together, these results demonstrate that only the stZF isoform binds well to the C/EBP-ATF site in the CHOP promoter and activates transcription.

### stZF promotes cell death possibly through CHOP induction

CHOP plays an important role in ER stress-induced apoptosis. Since stZF induces CHOP expression, we were interested in investigating whether stZF plays any role in apoptosis.

RAW264.7 cells were transfected with stZF or pcDNA3.1 (control) and treated with 5 μg/ml of tunicamycin. Results from Alamar Blue assay were plotted as graphs (Fig. 5A). Overexpression of stZF reduced cell viability at all time points. An independent-samples t-test was conducted to compare cells expressing stZF and control cells. Significant difference was found at 6-hour time point (*p*=0.049), 17-hour time point (*p*=0.00047) and 24-hour time point (*p*=0.027). It took 33 hours to reduce 50% of viable cells in control cells, but only 24 hours for cells expressing stZF. The most dramatic decrease in fluorescence intensity occurred between the 9-hour and 17-hour time point – about 35% from cells expressing stZF and 5% in cells transfected with pcDNA3.1. This could be due to the increased induction of CHOP by stZF after the cells exposed to tunicamycin for a prolonged period of time (Fig. 3A). DNA laddering assay also suggested that a higher number of cells committed apoptosis when overexpressing stZF after the 17-hour time point (Fig. 5C).

**Figure 5.**
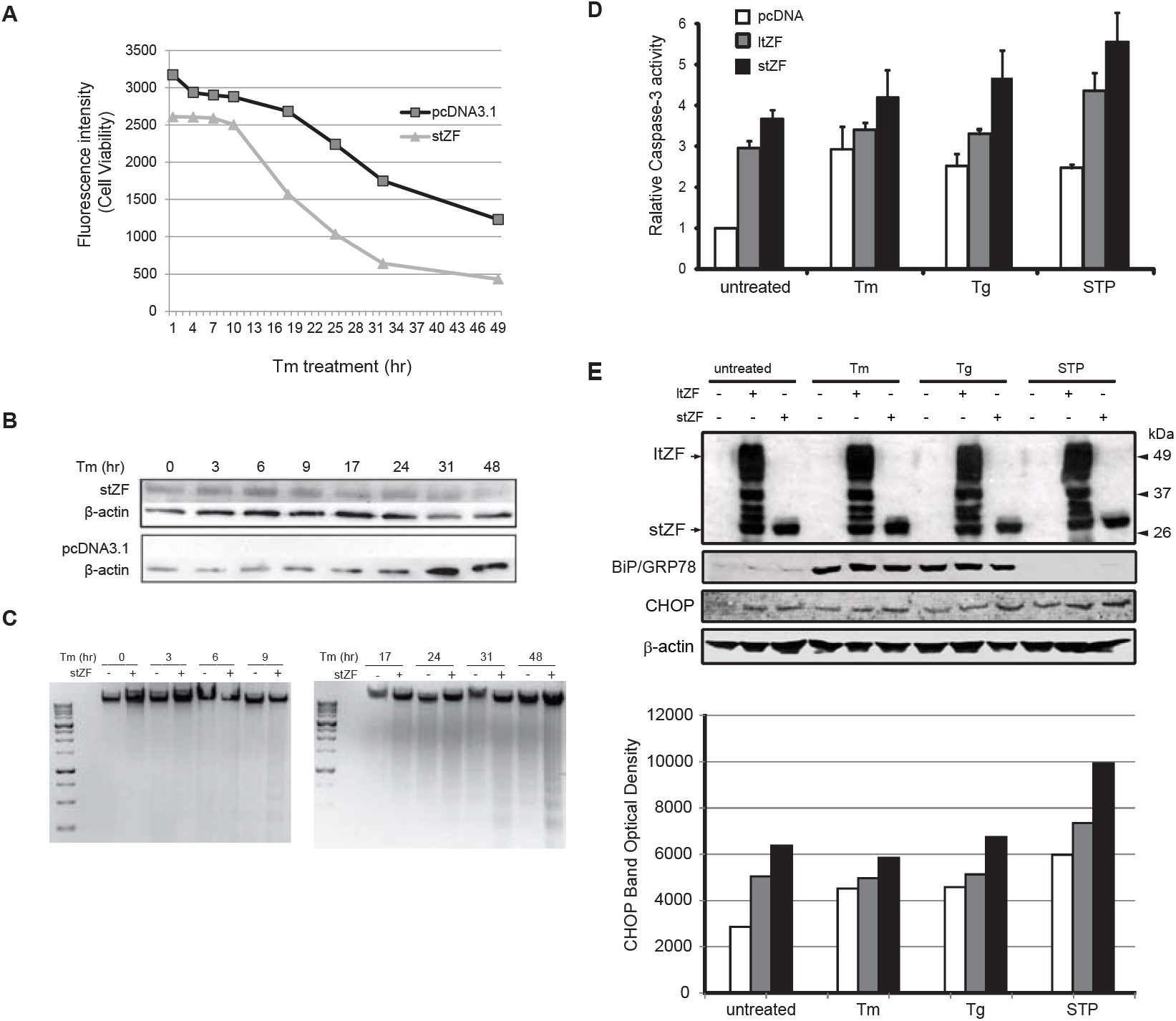
Overexpression of stZF reduces cell viability and induces apoptosis. (A) An Alamar Blue assay was performed to measure cell viability. RAW264.7 cells were transfected with stZF or pcDNA3.1 and treated with 5 μg/ml tunicamycin. Overexpression of stZF decreased cell viability (fluorescence intensity) at all time points of treatment. Significant difference in cell viability were detected in cells treated for 6 hours (*p*=0.049), 9 hours (*p*=0.00047) and 17 hours (*p*=0.027) by student’s t-test. (B) Cell lysates from the Alamar Blue assay were probed by 1486.4 (anti-ZF) and analyzed by western blots. The results confirmed the protein expression of stZF in cells transfected with stZF. Both blots (stZF and pcDNA3.1) were detected by the ChemiDoc™ XRS+ system (Bio-Rad) for same amount of time. (C) Overexpression of stZF induces DNA fragmentation. RAW264.7 cells transfected with stZF (labeled as “st”) showed higher amounts of fragmented DNA compared to cells transfected with pcDNA3.1 (labeled as “pc”), after they were exposed to tunicamycin for 17 hours or longer. Genomic DNA was collected from the transfected cell lysates and analyzed on an agarose gel. Three μg of DNA was loaded for each lane (quantified by Nanodrop8000). Both gels were detected together. Fragmentation of genomic DNA is a hallmark for late apoptosis. (D) Caspase assay of HeLa cells transfected with plasmids encoding ltZF and stZF isoforms. These cells were either untreated or treated with 2 μg/ml tunicamycin or 1 μM thapsigargin for 24 hours or 100 nM staurosporine (STP) for 5 hours prior to measuring caspase-3 activities. (E) Western blot analysis of the cell lysates from the caspase assays.

Caspase-3 activity was measured and used to monitor apoptosis. We found that overexpression of stZF strongly induced caspase-3 activity both in untreated cells (Fig. 5D) and in cells treated with ER stressors such as tunicamycin (2 μg/ml, 24 hrs) and thapsigargin (1 μM, 24 hrs), as well as in cells treated with apoptosis inducer staurosporine (100 nM, 5 hrs). Although ltZF was also able to stimulate apoptosis (except in tunicamycin-treated samples), it was found to be less effective than stZF (Fig. 5D). Similarly, cells overexpressing stZF were also found to have higher amount of DNA laddering, another hallmark of apoptotic cells (Fig. 5C). To investigate whether ltZF and stZF induced cell death correlates with the induction of endogenous CHOP, Western blot analysis were carried out using the same cell lysates used in the caspase-3 assays. The levels of the CHOP protein were found to be elevated in response to both stZF and ltZF. However, CHOP expression stimulated by stZF transfection was found to be significantly higher than that of ltZF, and corresponds to the levels of caspase activity (Fig. 5E).

Taken together, these results suggest that stZF stimulates caspase activation and apoptosis through CHOP expression.

### stZF is more stable under ER stress

Our results showed that protein of stZF can be induced by prolonged ER stress. We hypothesized that stZF is stable under ER stress, since its presence in late UPR is required to induce apoptosis. To analyze the stability of stZF, HEK293 cells were transfected with stZF and then treated with cycloheximide [72]. These cells were compared with those that were transfected with stZF but exposed to 5 μg/ml of tunicamycin for 24 hours prior to the cycloheximide treatment. Densitometric analysis was performed on western blots (Fig. 6). Cells were collected at the indicated time points post-cycloheximide treatment to analyze the half life of stZF. The results reveal that in the presence of ER stress, the half life of stZF is increased to 5 hrs, when compared to its half-life of 2 hrs in the absence of any ER stress (Fig. 6).

**Figure 6.**
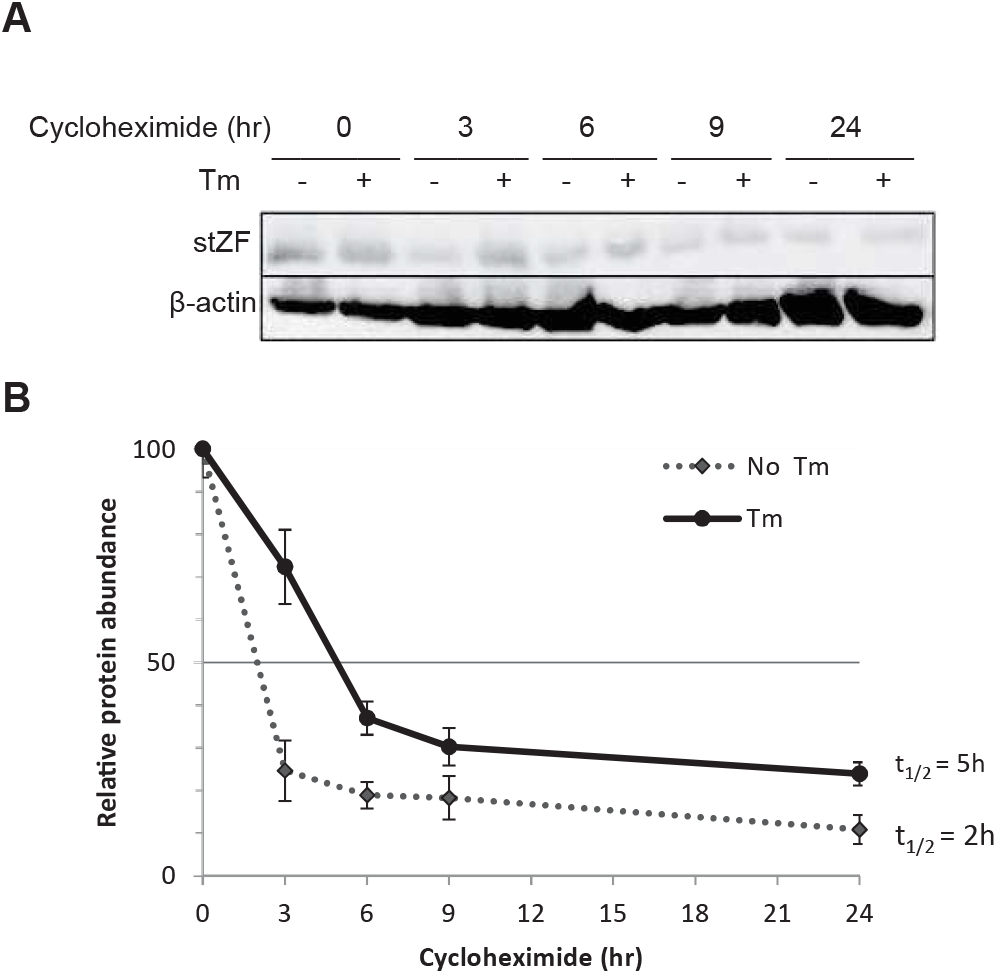
stZF is stable under prolonged ER stress. (A) HEK293 cells were transfected with stZF and half of the cells were treated with 5 μg/ml tunicamycin (indicated with “+”) for 24 hours prior to the cycloheximide treatment (50 μg/ml). Protein of stZF was more stable in cells treated with Tm. (B) Densitometric analysis for western blots of cycloheximide chase assay was performed by the software ImageJ. Relative protein levels at the zero hour time point were set to 100%. Protein expression was normalized to beta-actin levels. Protein half-life of stZF was 2.5 times longer when the cells were exposed to tunicamycin prior to the cycloheximide treatment. Results from two independent experiments are shown with standard errors.

## DISCUSSION

The UPR is primarily an adaptive response for the cell to adjust to stress and restore homeostasis in the ER. In response to sustained stress, the UPR shifts the balance from pro-survival to pro-cell death. In this report, we have investigated the production of 4 different isoforms of CREBZF. We have identified that a particular isoform of CREBZF called stZF, is specifically induced by prolonged ER stress through alternative translation initiation and mRNA splicing. This resultant stZF isoform is a potent transcription factor with a well characterized N-terminal transcription activation region (Cockram et al., 2006; Hogan et al., 2006; Lu and Misra, 2000b) and an IFFFR pentapeptide tail (Fig. 7). It activates CHOP transcription through a C/EBP-ATF element in the promoter and promotes apoptosis. As the nucleotide sequence in this gene region is perfectly conserved among several CREBZF homologs (Fig. 1C), we believe that the molecular mechanism for the functional induction of stZF may be a critical component of UPR signaling.

**Figure 7.**
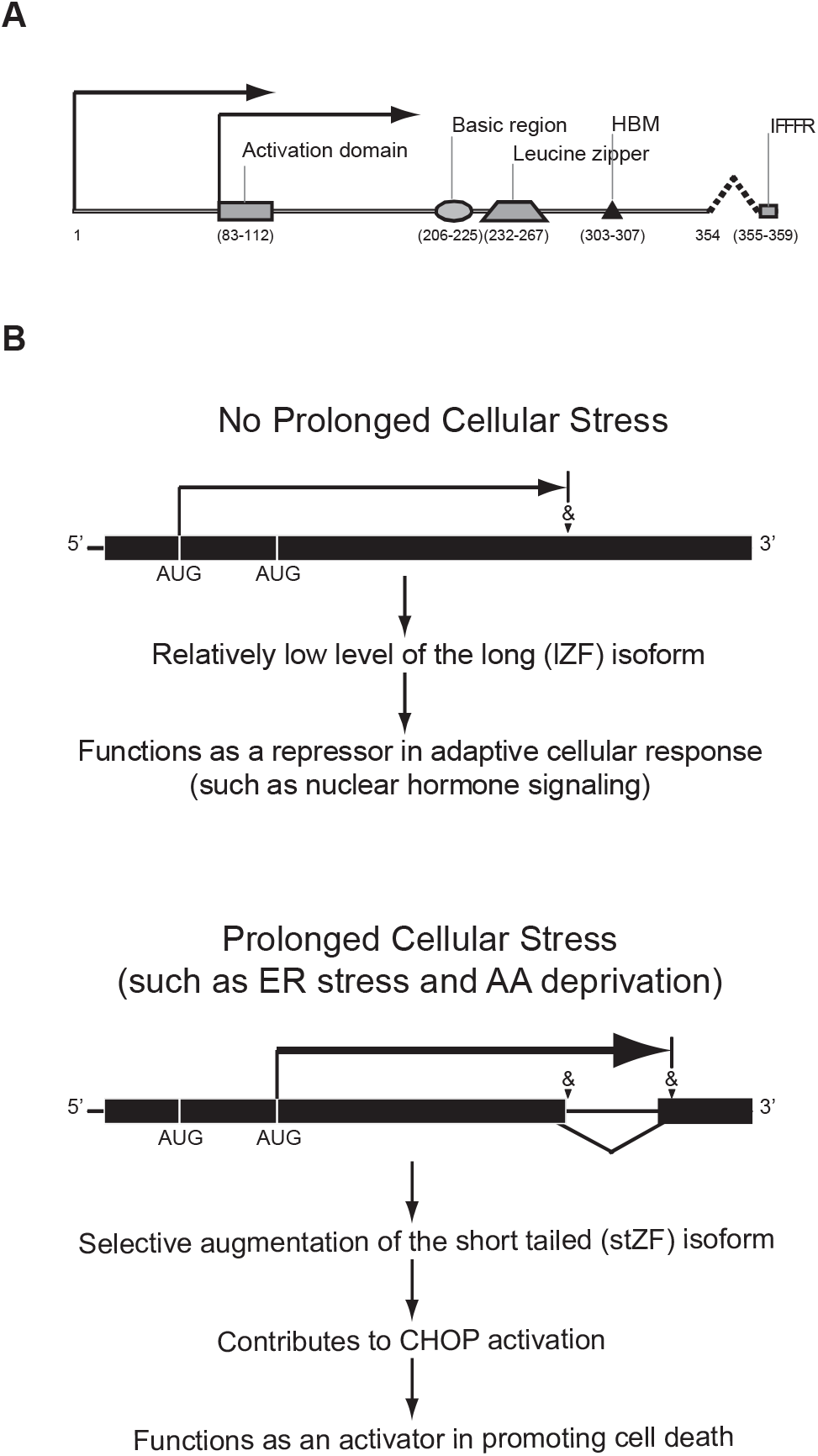
A working model of CREBZF isoform induction and their functions. (A) Schematic diagram of CREBZF protein depicting the known functional domains, as well as alternative translational initiation and mRNA splicing. (B) In cells without prolonged cellular stress, the CREBZF transcript by default is not spliced, and translation starts from the first 5’ AUG. The resultant lZF functions primarily as a repressor in an adaptive cellular response such as hormone signaling. In the event of prolonged cellular stress (such as ER stress or amino acid deprivation), CREBZF mRNA is alternatively spliced at the indicated site and protected from degradation by NMD. In addition, a downstream AUG is used to initiate translation. As a result, the stZF isoform with a potent N-terminal activation domain is produced, which contributes to CHOP induction and promotes cell death.

The original short non-tailed CREBZF (Lu and Misra, 2000b) was identified through a yeast two-hybrid screen using the cell cycle regulator protein HCF-1 (Julien and Herr, 2003, 2004; Tyagi et al., 2007) as bait. Although it has been known for some time that CREBZF is a transcription factor, its downstream targets have been elusive. Because of the unique basic region that is significantly different from other bZIP proteins, CREBZF is not believed to be able to bind to DNA by itself as a homodimer. Instead, CREBZF exerts its function as a transcription factor by associating with and modifying the action of other DNA-binding transcription factors (Cockram et al., 2006; Hogan et al., 2006; Lu and Misra, 2000b). In this regard, CREBZF is similar to CHOP (Oyadomari and Mori, 2003; Ron and Habener, 1992; Ubeda et al., 1996; Weidenfeld-Baranboim et al., 2008) in that, dependent upon the context of protein-promoter complexes, CREBZF can function both as a positive and a negative regulator of other transcription factors (Akhova et al., 2005; Hogan et al., 2006; Misra et al., 2005; Valderrama et al., 2008; Xie et al., 2008; Xie et al., 2009a). Interestingly, the current study indicates that CHOP is also a downstream target of CREBZF in signaling cell death during prolonged cellular stress. CREBZF appears to be widely expressed, primarily in the form of the long non-tailed isoform lZF (Lu and Misra, 2000b; Xie et al., 2008), while sZF was detected in HeLa cells under non-stress conditions (Xie et al., 2008).

Two major events are necessary during the UPR for the cell to shift from lZF to stZF protein production: 1) alternative translation initiation at the second AUG; and 2) alternative splicing and preferential protection of the mRNA variant from NMD to generate the pentapeptide IFFFR tail. It is well known that, in mammalian systems, the −3 and +4 positions of the mRNA are important for ribosome recognition, with RnnAUGG (R is a purine) being the optimal context and YnnAUGH (Y, pyrimidine; H, not a “G”) being most leaky (Kozak, 2005). If the first start codon in the coding sequence is in a suboptimal context, translation may be initiated at a downstream AUG by the ribosomal leaky scanning mechanism. For CREBZF, however, despite optimal initiation context at the second AUG (Fig. 1B), the short CREBZF appears to be produced primarily under prolonged stress conditions. A similar observation has also been made with another protein, C/EBPα/β, where a downstream AUG is preferentially used under cellular stress (Ariyama et al., 2008; Calkhoven et al.). Like CREBZF, often these truncated polypeptides translated from a downstream AUG have different cellular functions, as in the case of human glucocorticoid receptor α (Gu et al., 2008; Yudt and Cidlowski, 2001), MDM2 (Cheng and Cohen, 2007) and p53 (Courtois et al., 2002). For the tailed isoforms of CREBZF to be made the cell also has to overcome the targeted mRNA degradation of the alternative splicing variants by NMD. It is known that eukaryotic cells use NMD to regulate gene expression (Matsuda et al., 2008; Stalder and Muhlemann, 2008). In the case of CREBZF, however, it is remarkable to observe multiple mechanisms that work in concert to produce the stZF isoform that is a potent transcription activator of CHOP. Clearly more research is necessary to elucidate how the prolonged ER stress signal is converted to control translation initiation, mRNA splicing and NMD regulation of CREBZF.

In this report, we found that only the tailed isoforms of CREBZF could activate CHOP expression through the C/EBP-ATF element. However, the mechanism for the functional significance of the pentapeptide IFFFR at their C-terminus still remains unclear. Since the non-tailed CREBZF isoforms, lZF and sZF, are already located in the nucleus (Lu and Misra, 2000b; Xie et al., 2008), the pentapeptide addition to the CREBZF protein is unlikely to serve as a subcellular targeting signal for its transcription factor function. The presence of three bulky phenylalanine residues and one positively charged arginine may indicate that it is a protein binding motif. Synthetic pentapeptide IFFFR has been reported to bind TNF-α and inhibit TNF-α from binding its receptors (Kruszynski et al., 1999). The unique IFFFR tail may also change the conformation of the polypeptide and therefore affects post-translational modification of adjacent residues, for instance, blocking the phosphorylation of the adjacent serine residue (Fig. 1C). The study of impact of the pentapeptide tail on protein modification is currently under way.

Although the ltZF-M83I mutant that cannot produce stZF still activated transcription from the C/EBP-ATF element, its activation potential was significantly weaker than that of stZF (Fig. 4B). The benefit of translating from the downstream AUG may be the transactivation potential: the N-terminal activation domain in sZF appeared more potent compared to when it is embedded behind the 82 amino acids in lZF, as seen with the CHOP promoter (Fig. 3A) or when tethered to a GAL4 promoter (data not shown).

In the mammalian UPR, CHOP induction is primarily mediated through the PERK/eIF2α/ATF4 UPR branch, although the IRE1α/XBP1 and ATF6α pathways also contribute (Acosta-Alvear et al., 2007; Fawcett et al., 1999; Harding et al., 2000; Wu et al., 2007; Yamamoto et al., 2007). ER stress activated PERK phosphorylates eIF2α, which leads to attenuation of global translation but selective enhancement of ATF4 and induction of CHOP. In addition to PERK, GCN2 is one of several other kinases that also phosphorylate eIF2α to activate a downstream gene expression program that is referred to as the integrated stress response (ISR) (Harding et al., 2003), which includes ATF4 and CHOP. GCN2 specifically responds to amino acid deprivation (Guo and Cavener, 2007; Hao et al., 2005; Maurin et al., 2005); and the downstream target CHOP has been found to be one of the key gene regulators in the transcription factor network during the amino acid stress response (Bruhat et al., 1997; Jousse et al., 1999; Su and Kilberg, 2008).

It is worth noting that both ATF4 (Lu et al., 2004; Vattem and Wek, 2004) and CHOP (Jousse et al., 2001) are also regulated at the post-transcriptional level. Instead of upstream AUGs as in the case of CREBZF, the mRNA of both genes contains upstream Open Reading Frames (uORFs), which repress translation of the downstream main ORF. Translation of ATF4 and CHOP is enhanced by a ribosomal leaky scanning and/or reinitiation mechanism during the cellular response to ER stress or amino acid deprivation, which bears remarkable similarities with CREBZF.

In agreement with CHOP, the stZF is also found to be induced by amino acid deprivation (Zhang et al., 2010). Furthermore, while CHOP is generally considered as a pro-apoptotic factor which is induced prior to apoptosis, and suppression of CHOP expression prevents apoptosis (Marciniak et al., 2004; Oyadomari et al., 2001; Zinszner et al., 1998), we found that overexpression of stZF strongly induces apoptosis as measured by caspase-3 activity which is enhanced by 4-fold, even in the absence of any stress treatments (Fig. 5D). Overexpression of stZF also increased apoptotic DNA fragmentation (Fig. 5C). Cellular metabolism is also reduced by stZF overexpression (Fig. 5A), especially at the time point that stZF induced protein level of CHOP.

Based on these observations, we propose that 1) CREBZF is a key component of the integrated stress response signaling pathway; 2) while the non-tailed isoforms of CREBZF (i.e., lZF and sZF) may function as transcriptional repressors in the nuclear receptor signaling, the tailed isoform (or the stZF specifically) which is tightly regulated and induced primarily late in the cellular stress response is essential for the role of CREBZF in inducing CHOP and promoting cell death during the late stage of the UPR (Fig. 7). Since CHOP and ISR have been implicated in many diseases such as cancer, diabetes and neurodegenerative diseases, it should be rewarding to further investigate the role of CREBZF in the context of interactions with other members of this pathway.

## METHODS AND MATERIALS

### DNA constructs

The following CHOP promoter reporter plasmids were described previously (Bruhat et al., 2000), with numbers in parentheses denoting nucleotide positions of the start and end of the promoter fragments: pGL3-CHOP (−954/+91) containing the wild type CHOP promoter; the reporter plasmids containing progressive 5’ deletions of the CHOP promoter pGL3-CHOP (−649/+91), pGL3-CHOP (−249/+91), pGL3-CHOP (−221/+91), pGL3-CHOP (−119/+91), pGL3-CHOP (−40/+91); the reporter plasmid pGL3-CHOP (−649/+91) containing the disrupted SP-1 element and pGL3-CHOP (−649/+91) containing the disrupted C/EBP-ATF element; and the reporter plasmid containing 2 copies of (C/EBP)-ATF elements 5’-ATTGCATCAT. The pRL-SV40 plasmid (Promega) contains the Renilla Reniformis luciferase gene under control of the SV40 immediate-early promoter. sZF (pcZF, the original Zhangfei) was described previously (Cockram et al., 2006; Hogan et al., 2006; Lu and Misra, 2000b). The cDNAs encoding lZF, ltZF and stZF (Fig. 1A) were generated by RT-PCR and cloned into the EcoRI/XhoI site of pcDNA3.1 (Invitrogen) to create lZF, ltZF and stZF. QuikChange site-directed mutagenesis (Stratagene) was used to create lZF-M83I and ltZF-M83I from the wild type lZF and ltZF plasmid, respectively, in which the 83rd amino acid of lZF and ltZF protein was changed from methionine to isoleucine by substituting the ATC for ATG using the following primers: 5’-CTCCCCCGAGGAG**ATC**GAGGAGGAGGCG, and 5’-CGCCTCCTCCTC**GAT**CTCCTCGGGGGAG (the mutated 83rd amino acid-codon is in bold).

### Cell culture, transfection and ER stress treatment

HeLa and human embryonic kidney (HEK) 293 cells were cultured in Dulbecco’s Modified Eagle’s medium (high glucose, Sigma) supplemented with 10% (v/v) fetal bovine serum (Invitrogen), 100 IU/ml penicillin and 100 μg/ml streptomycin. Mouse embryonic fibroblast (MEF) and RAW264.7 (mouse macrophage) cells were cultured in Dulbecco’s Modified Eagle Medium (high glucose; Invitrogen) supplemented with 10% (v/v) fetal bovine serum (Invitrogen), 1% (v/v) penicillin/streptomycin (Lonza), 1% MEM Non-essential Amino Acid Solution (100×) (Sigma-Aldrich) and 0.1% Beta-Mercaptoethanol (Fisher Scientific). All cultures were maintained in a 5% CO_2_ humidified atmosphere at 37 °C and passaged every 2-3 days. Cells were grown to approximately to 70% confluency prior to transfection using the calcium phosphate precipitation method (Lu and Misra, 2000a) for luciferase assays, and DreamFect (OZ Biosciences) for caspase-3 assays. For transfection using FuGENE® HD (Roche), cells were seeded into six-well plates, 96-well plates, 6-cm dishes or 10-cm dishes with 100ul, 2ml, 5ml or 10ml of growth media respectively. Cells were grown to 75% confluency prior to the transfection. Media was changed 12 hours post-transfection. In the case of transfection using Xfect™ (Clontech), cells were grown to 50% confluency prior to the transfection. Media was changed 4 hours (HeLa and HEK293 cells) or 8 hours (MEF and RAW264.7 cells) post-transfection. Cells were allowed to grow until they reached 80-85% confluency before exposing them to tunicamycin (Sigma-Aldrich). All cells (untreated and treated) were harvested at the end of treatment.

### Total RNA isolation, RT-PCR and Southern blotting

Total RNA was isolated with Trizol (Invitrogen) from cultured HeLa cells. SuperScriptII reverse transcriptase (Invitrogen) and primer 5’-AAAAACAGCAGAAGCCCTGA were used to synthesize the cDNA. The primers used in the reverse transcription-PCR (RT-PCR) were the following: 5’-ACTGGACTGGCTCGCTTG and 5’-CAGCTGGTCCCGTCTCTTCT. After electrophoresis, the agarose gel was transferred to a Hybond-N+ nylon membrane (GE Healthcare). An 819-bp fragment containing the full length sZF cDNA was labeled by random priming with [α-^32^P] dCTP and used as a probe.

### Western blotting and immunoprecipitation

Detection of CREBZF was performed using a polyclonal antibody (Rb6) raised against the four isoforms of CREBZF. Other primary antibodies used include: a polyclonal CHOP antibody (Santa Cruz), a polyclonal BiP/GRP78 antibody (Santa Cruz Biotech) and a β-actin monoclonal antibody (clone AC-15; Sigma). Blots were visualized using ECL Plus (GE Healthcare) on a Typhoon 9400 Phosphorimager (GE Healthcare). For the detection of endogenous ZF in RAW264.7 cells, immunoprecipitation was performed by using Dynabeads® Protein G beads (Invitrogen) as per manufacturer’s instructions. 105 micrograms of total protein were added to the beads, protein concentration was quantified by BSA Protein Assay. Two micrograms of monoclonal anti-ZF (mouse) was used in the immunoprecipitation, and polyclonal 1486.4 antibody (rabbit) was used in western blots.

### Cell Viability Assay

HeLa cells were plated at a concentration of 10,000 cells/well in a 96-well plate with 200 μl of media, and allowed to adhere and recover for 18 hours prior to treatment. Cells were then treated at 12-hour intervals for 48 hours with 2 μg/ml tunicamycin. At 4 hours prior to the final measurement, 100 μl of resazurin (50 μM, Sigma #R7017) was added to each well. The plate was read immediately on a Biotek FLX100 fluorescence microplate reader (Ex516/20, Em590/35) to correct for background, and then returned to the incubator for 4 hours before a second reading. Two biological repeats were performed in triplicates for each treatment. Averages of the percentage of stained cells compared to non-treated cells were plotted as percent of viability with standard errors. The same procedures were followed for RAW264.7 cells, but the length of treatment time varied and 5 μg/ml tunicamycin was used. Results were analyzed by independent-samples (student’s) t-test. A significance level of 0.05 was used.

### Cycloheximide (CHX) chase assay

HEK 293 cells were transiently transfected. At 24 hour post-transfection, cells were treated with 5ug/ml of tunicamycin (Tm). Twenty-four hours after the addition of Tm, fifty percent of cells were treated with 50ug/ml of cycloheximide (Sigma-Aldrich) for different lengths of time. Untreated and treated cells were harvested at the same time by the trypsinization.

### Measurement of caspase-3 activity

Cell were lysed with a lysis buffer containing 50 mM HEPES [pH 7.4], 0.1% 3-[(3-cholamidopropyl)-dimethylammonio]-1-propanesulfonate [CHAPS], 0.1 mM EDTA. Lysates were incubated with the caspase-3 substrate DEVD-AMC [N-acetyl-Asp-Glu-Val-Asp-(7-amino-4-methylcoumarin); Biomol Research Laboratories]. The continuous liberation of AMC was examined at 37°C using a Bio-Tek FLx800 microplate fluorescence reader (Bio-Tek, Winooski, VT) with an excitation wavelength of 380 nm and emission at 460 nm. The fluorescence units of AMC released/min/μg protein were calculated for all the samples. Assays were repeated independently at least three times, and results are shown with standard errors.

### Dual luciferase assays

Cells were co-transfected with 1.0 μg/35-mm dish of the effector plasmids and 1.0 μg/dish of the pGL3-CHOP luciferase reporter plasmids or pGL3-Basic (Promega) with a simian virus 40 (SV40) promoter, together with 40ng/well of Renilla luciferase plasmid pRL-SV40 (Promega) as an internal control. At 24 hours post transfection, media was changed and the cells were allowed to recover for 24 hours in untreated or 2 μg/ml tunicamycin-treated conditions. Cells were harvested and lysed, and dual luciferase assays were performed according to the manufacturer’s protocols (Promega). Luciferase activity was measured using a Turner ED-20e Luminometer and calculated as relative luciferase activity (firefly luciferase/Renilla luciferase) to correct for transfection efficiency. Assays were repeated independently at least three times, and results are shown with standard errors.

### Chromatin Immunoprecipitation (ChIP)

HeLa cells cultured in 10-cm plates were transfected with 5 μg/plate of pcFLAG, pcFLAG-sZF, pcFLAG-lZF, pcFLAG-stZF, pcFLAG-ltZF, pcFLAG-lZF-M83I, pcFLAG-ltZF-M83I along with CHOP promoter as described above, respectively. The ChIP assay was performed by following the protocol reported in (Zhou et al., 2004) with minor modifications. Briefly, after cross-linking in 1% formaldehyde, the cells were lysed and sonicated. The supernatant was pre-cleared with Protein A beads (Amersham). Equal amounts of samples were used in the immunoprecipitation. A 5% aliquot of the pre-cleared chromatin was taken as input, and the rest was incubated with 2 μg of M2 anti-FLAG monoclonal antibody (Sigma) or 2 μg of H3 polyclonal antibody (FL-136; Santa Cruz Biotechnology). After reversing the formaldehyde-induced cross-linking, the chromatin DNA was used in PCR to produce a 216-bp CHOP promoter product with the primers 5’-GGCGACCAAGGCTGATAG and 5’-AGGCTTCACGGAGGAGGT. Similarly, primers 5’-TCCTTCCCAGCCAGAAGTTT and 5’-TCCCACTAGCCTCACCTCTG for the MCM5 promoter were used in the control ChIP assays. The gel electrophoresis images were acquired on a Bio-Rad Gel Doc 2000 gel documentation system after ethidium bromide staining.

## ACKNOWLEDGEMENTS

We thank Pierre Fafournoux for providing the pGL-CHOP plasmids, Richard Mosser for help with the Alamar Blue and caspase-3 assays, Yaping Jin for technical support, and Cheryl Cragg and Adam McCluggage for helpful discussions. This work was supported by a Natural Sciences and Engineering Research Council Discovery Grant awarded to R.L.; and the Scholarship Council of Chinese Ministry of Education to Y.Z.

## Author Contributions

YL, WKY and TT performed experiments, analyzed data and wrote the manuscripts; YZ, MZ, and TEA designed and initiated experiments; RL supervised all experiments and data analysis, and wrote manuscript.

## Abbreviations

C/EBP: CCAAT enhancer binding protein
ER: endoplasmic reticulum
UPR: unfolded protein response
PERK: protein kinase-like endoplasmic reticulum kinase
ATF6: activating transcription factor 6
IRE1: endoribonuclease inositol-requiring enzyme-1
ATF4: activating transcription factor 4
ERAD: ER-associated degradation
HSV-1: Herpes Simplex Virus-1
bZIP: basic region leucine zipper
ERSE: ER-stress response element
lZF: long CREBZF isoform
sZF: short CREBZF isoform
ltZF: long-tailed CREBZF isoform
stZF: short-tailed CREBZF isoform
NMD: nonsense-mediated mRNA decay
ChIP: Chromatin Immunoprecipitation
RT-PCR: Reverse transcriptase PCR
ISR: integrated stress response.

